# Stochastic models of cell invasion with fluorescent cell cycle indicators

**DOI:** 10.1101/273995

**Authors:** Matthew J Simpson, Wang Jin, Sean T Vittadello, Tamara A Tambyah, Jacob M Ryan, Gency Gunasingh, Nikolas K Haass, Scott W McCue

**Affiliations:** School of Mathematical Sciences, Queensland University of Technology, Brisbane, Australia; The University of Queensland, The University of Queensland Diamantina Institute, Translational Research Institute, 37 Kent St, Woolloongabba, Brisbane, QLD 4102, Australia; Discipline of Dermatology, Faculty of Medicine, Central Clinical School, University of Sydney, Sydney, NSW, Australia

**Keywords:** cell invasion, cell migration, cell proliferation, cell cycle, stochastic model, partial differential equation

## Abstract

Fluorescent cell cycle labelling in cell biology experiments provides real time information about the location of individual cells, as well as the phase of the cell cycle of individual cells. We develop a stochastic, lattice-based random walk model of a two-dimensional scratch assay where the total population is composed of three distinct subpopulations which we visualise as red, yellow and green subpopulations. Our model mimics FUCCI technology in which cells in the G1 phase of the cell cycle fluoresce red, cells in the early S phase fluoresce yellow, and cells in the S/G2/M phase fluoresce green. The model is an exclusion process so that any potential motility or proliferation event that would place an agent on an occupied lattice site is aborted. Using experimental images and previous experimental measurements, we explain how to apply the stochastic model to simulate a scratch assay initialised with a low to moderate density monolayer of human melanoma cell line. We obtain additional mathematical insight by deriving an approximate partial differential equation (PDE) description of the stochastic model, leading to a novel system of three coupled nonlinear reaction diffusion equations. Comparing averaged simulation data with the solution of the continuum limit model confirms that the PDE description is accurate for biologically-relevant parameter combinations.

## 1. Introduction

Cell invasion involves moving fronts of populations of cells. The movement of these fronts is driven by combined cell migration and cell proliferation [1]. Such moving fronts play an important role in tissue repair [2, 3], malignant spreading [4, 5, 7] and embryonic development [8, 9]. Cell invasion is most commonly studied *in vitro* using a scratch assay [10, 11], where a monolayer of cells is grown on a two-dimensional substrate and a portion of the population is scratched away using a sharp-tipped instrument. After the scratch has been made, individual cells in the population migrate and proliferate, with the net result being the formation of a moving front of cells that closes the scratch [10, 11].

Cell invasion involves combined cell migration and cell proliferation. Cell proliferation is often conceptualised as a sequence of four phases: gap 1 (G1); synthesis (S); gap 2 (G2); and the mitotic (M) phase [12]. Together, the G1, S and G2 phases are referred to as *interphase*, which involves cells undergoing preparation for division. Following interphase, cells enter the mitotic phase, during which they divide into two daughter cells. Traditional scratch assays provide no means of distinguishing between different cell phases within the population. However, in 2008, fluorescent ubiquitination-based cell cycle indicator (FUCCI) technology [13] was developed and this technology allows us to visualise the cell cycle for individual cells within a population of cells [12–16]. FUCCI involves two fluorescent probes in the cell which emit red fluorescence when the cell is in the G1 phase, or green fluorescence when the cell is in the S/G2/M phase. FUCCI is important because it allows us to visualise cell cycle transitions in real time. FUCCI technology has many important applications, including developmental biology and cancer biology [17, 18]. For example, in cancer biology, many important drugs involve interrupting the cell cycle [19, 20]. Therefore, it is useful to be able to visualise, in space and time, how cells progress through the cell cycle.

In this work we describe a new stochastic random walk model of cell migration and cell proliferation that can be used to mimic, interpret and design scratch assay experiments performed with FUCCI technology. We begin by presenting images from a recent scratch assay performed with a human melanoma cell line. In these experiments, cells have been transfected with FUCCI so that each cell fluoresces either red, yellow or green [16]. Images in Figures 1(a)–(b) show a population of 1205Lu melanoma cells, with each cell fluorescing either red, yellow or green. The image in Figure 1(a) corresponds to *t* = 0 h, just after the scratch has been made in the uniform monolayer of cells. The image in Figure 1(b) shows the same field of view at *t* = 48 h, after the wound has essentially closed. To describe this kind of data, we propose a lattice-based random walk model that treats the total population of cells as three interacting subpopulations: (i) a subpopulation of red agents; (ii) a subpopulation of yellow agents; and (iii) a subpopulation of green agents. Each agent in each subpopulation undergoes a nearest neighbour random walk at a particular rate [21] and is permitted to progress through the cell cycle as red agents transition to yellow agents, and yellow agents transition to green agents, before green agents eventually divide into two red daughter agents. The final transition from green agents into two red agents is permitted only when there is sufficient space available. A schematic of this cell cycle progression is shown in Figure 1(c). Since the random walk model is an exclusion process [22], each lattice site cannot be occupied by more than one agent. Therefore, potential motility events or potential transition events that would place an agent onto an occupied lattice site are aborted [21, 23, 24]. This means that the model incorporates both contact inhibition of migration and contact inhibition of proliferation, which are both known to be important in two-dimensional scratch assays [25].

**Figure 1:**
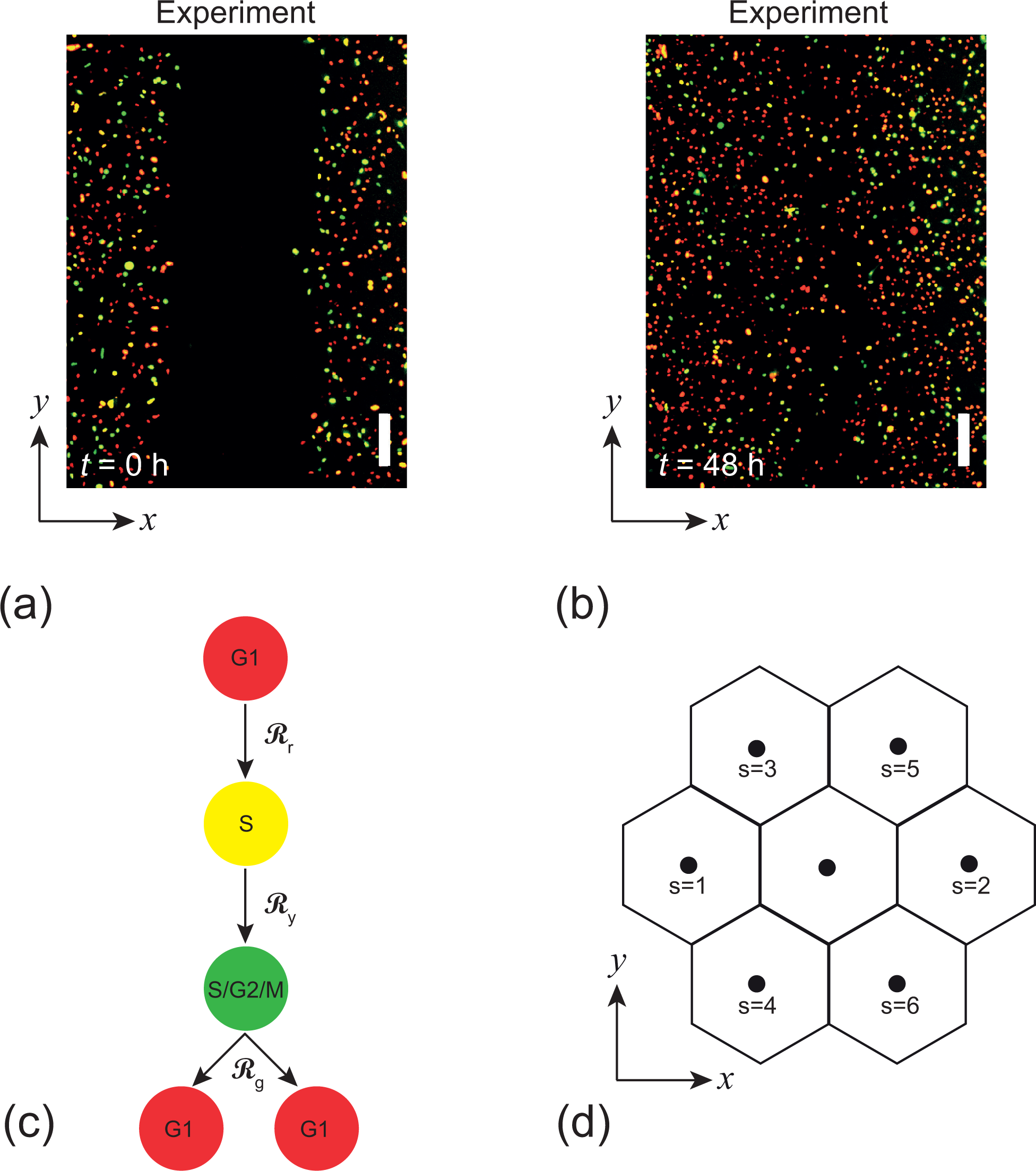
(a)–(b) Scratch assay with 1205Lu melanoma cells transfected with FUCCI. Images are shown at *t* = 0 h and *t* = 48 h [16]. Each scale bar corresponds to 200 µm. (c) Schematic showing the progression through the G1 phase (red), early S phase (yellow) and S/G2/M phase (green) for FUCCI. (d) Nomenclature for the hexagonal lattice, with lattice spacing Δ. Thevcentral lattice site, which we denote as the *k*^th^ site, is at location (*x, y*). Th e six nearest neighbour sites are at 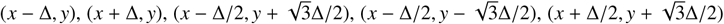 and 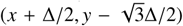, which we denote *s* = 1, 2, 3, 4, 5 and 6, as indicated in (d). Parameters in the discrete model include the lattice spacing Δ, the movement rates *M*_*r*_, *M*_*g*_ and *M*_*y*_, and the transition rates ℛ_*r*_, ℛ_*g*_ and ℛ_*y*_. The parameters in the continuum limit model are given by *D*_*r*_ = lim_Δ→0_(*M*_*r*_Δ^2^)/4, *D*_*y*_ = lim_Δ→0_(*M*_*y*_Δ^2^)/4 and *D*_*g*_ = lim_Δ→0_(*M*_*g*_Δ^2^)/4.

In addition to describing the stochastic model, we also derive a mean field continuum limit description of the stochastic model. This kind of approach has been pursued in many previous studies, with both single-species latticebased stochastic models (including [26, 27] for example) and single-species lattice-free stochastic models [28–32]. In contrast, the experimental system we consider here leads us to treat the population of agents in the discrete model as three subpopulations. Therefore, the continuum limit description is a novel system of three coupled nonlinear partial differential equations (PDE). This approach of having both an individual-level stochastic description and a populationlevel continuum description of the same underlying process is useful for interpreting cell biology experiments as some experimental data are most naturally interpreted using population-level measurements and properties, while other experimental data from the same experiment are best interpreted in a stochastic, individual-based framework [33–37]. To explore how well the continuum limit PDE describes the stochastic model, we generate a suite of identically prepared realisations of the stochastic model and explore how well the average behaviour of the stochastic model is accurately described by the solution of the system of PDEs. We conclude by presenting some parameter sensitivity of the continuum-discrete match, and outline how the new models presented here can be extended in future studies.

This work is an extension of our previous work which focuses exclusively on the heuristic development of continuum models to simulate collective cell migration with FUCCI technology [16]. However, we note that working with continuum models alone provides no opportunity to generate insight into cellular-level behaviours, and continuum models cannot produce images and snapshots that are directly comparable with experimental data. Therefore, the current work aims to develop a discrete model of scratch assays with FUCCI technology that can produce data that is directly comparable with experimental images. Furthermore, we analyse the discrete model and show that the continuum limit description leads to a novel system of PDEs which is analogous to that studied in [16]. An interesting feature of this system is that the diffusion terms are nonlinear. This nonstandard nonlinearity arises as a consequence of the multispecies nature of the discrete system.

## 2. Results and Discussion

### 2.1. Discrete model

We formulate a discrete exclusion process-based random walk model on a hexagonal lattice which is based on previous studies that model cell proliferation and migration in two dimensions [5]. Each lattice site is indexed *k* ∈ [1, *K*], and is associated with a unique Cartesian coordinate, (*x, y*). The lattice spacing is Δ > 0, and each lattice site is associated with a set of six nearest neighbouring lattice sites, which we index *s* = 1, 2, …, 6, as indicated in Figure 1(d). The total population of agents is composed of three distinct subpopulations: red coloured agents represent cells in the G1 phase of the cell cycle; yellow coloured agents represent cells in the early S phase of the cell cycle; and green agents represent cells in the S/G2/M phase of the cell cycle. Each subpopulation undergoes an unbiased random walk, with exclusion, and agents from each subpopulation can transition to one other subpopulation to mimic the progression of the cell cycle. While we have chosen to use a hexagonal lattice, it would also be reasonable to use a square lattice, and previous studies have compared these different approaches for similar models [6].

We simulate the stochastic model using the Gillespie algorithm [38], in which two kinds of events take place: (i) migration events; and (ii) cell cycle transition events. In this work we make the usual assumption that the time between different events is exponentially distributed [3, 5, 36]. To simulate migration, we allow agents in the red, yellow and green subpopulations to undergo an unbiased nearest neighbour random walk with rates *M*_*r*_, *M*_*y*_ and *M*_*g*_, respectively. Any agent attempting to undergo a migration event will first randomly choose a target site from one of the six nearest neighbour lattice sites. Potential motility events will only proceed if the randomly chosen target site is vacant. This kind of exclusion process is a simple way to represent contact inhibition of migration.

To simulate transitions through the cell cycle, we allow red agents to transition into yellow agents at a rate ℛ_*r*_, and yellow agents to transition into green agents at a rate ℛ_*y*_, as indicated in Figure 1(c). We take a simple approach by supposing that both the red-to-yellow and the yellow-to-green transitions are unaffected by crowding. Green agents are allowed to transition into red agents at rate ℛ_*g*_, and we note that this transition depends on the availability of space to accommodate the new daughter agent, as indicated in Figure 1(c). For the green-to-red transition to take place, the occupancy status of a randomly chosen nearest neighbour lattice site surrounding the green agent is assessed. If the target site is vacant, the green agent transitions to red, and a new red daughter agent is placed on the target site.

There are many options for how we could treat a green-to-red transition where the target site is occupied. Here, we take the simplest possible approach and abort the event, leaving the green agent unchanged. Other options for modelling this situation might be to abort the division event but to allow the green agent to transition to red without dividing, or to abort the division event and remove the green agent from the lattice. Another option is to assess the availability of space at the beginning of the cell cycle, when the agent is red, and only allow that agent to progress through the cell cycle if there is sufficient space at that time. This option could be simulated by treating the red subpopulation as being composed of two additional subpopulations: (i) those red agents that have committed to progressing through the cell cycle; and (ii) those that are waiting for space to become available before they are able to progress through the cell cycle. This approach would require the consideration of four subpopulations in total and so we do not take this approach here. These and other potential variations could be incorporated into our stochastic framework, and different choices about the implementation of the discrete model will lead to different PDE descriptions. Regardless of the details of these choices, the same framework can be used to derive the corresponding PDE description.

### 2.2. Continuum models

In any one realisation of the discrete model the occupancy status of each site at time *t* can be represented as **R**_*k*_(*t*) = 1 if site *k* is occupied by a red agent, and **R**_*k*_(*t*) = 0 if site *k* is not occupied by a red agent. Similarly, **Y**_*k*_(*t*) = 1 if site *k* is occupied by a yellow agent, and **Y**_*k*_(*t*) = 0 if site *k* is not occupied by a yellow agent, and **G**_*k*_(*t*) = 1 if site *k* is occupied by a green agent, and **G**_*k*_(*t*) = 0 if site *k* is not occupied by a green agent. If we consider an ensemble of *N* identically prepared realisations of the stochastic model we can construct averages of these binary occupancies:

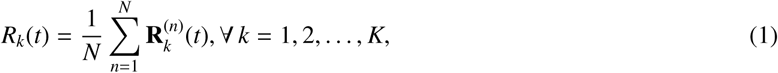

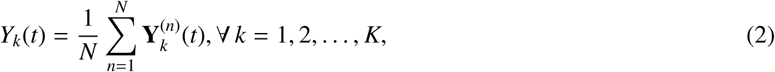

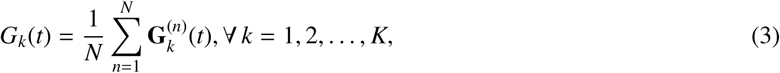

where 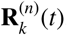, 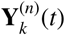 and 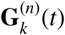 indicate the occupancy of site *k*, at time *t*, by a red, yellow and green agent in the *n*^th^ identically prepared realisation, respectively. After constructing this average we have *R*_*k*_(*t*) ∈ [0, 1], *Y*_*k*_(*t*) ∈ [0, 1] and *G*_*k*_(*t*) ∈ [0, 1], and we can interpret these averaged quantities as the probability that site *k* is occupied by a red, green or yellow agent at time *t*, respectively. It is also convenient to define a variable that describes the probability that site *k* is occupied, regardless of the subpopulation, *T*_*k*_(*t*) = *R*_*k*_(*t*) + *Y*_*k*_(*t*) + *G*_*k*_(*t*), noting that *T*_*k*_(*t*) ∈ [0, 1]. With these definitions we can formulate approximate conservation statements describing the time rate of change of occupancy of site *k* for the three different subpopulations [21, 23]

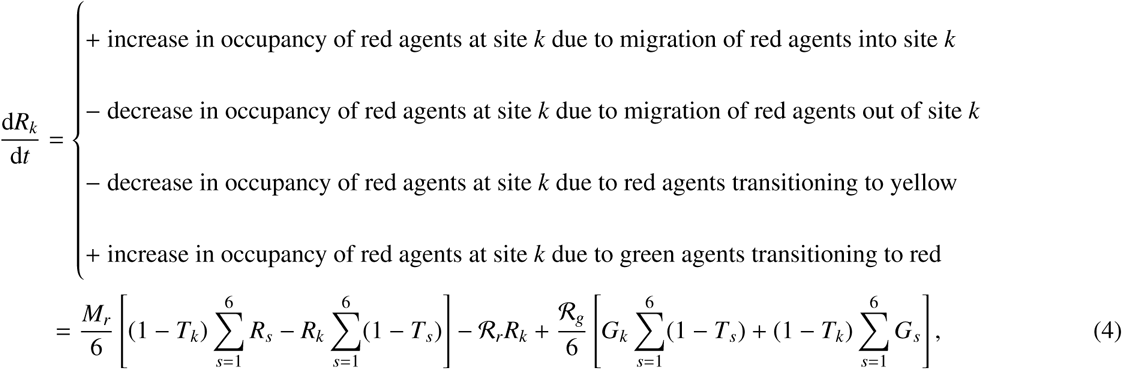

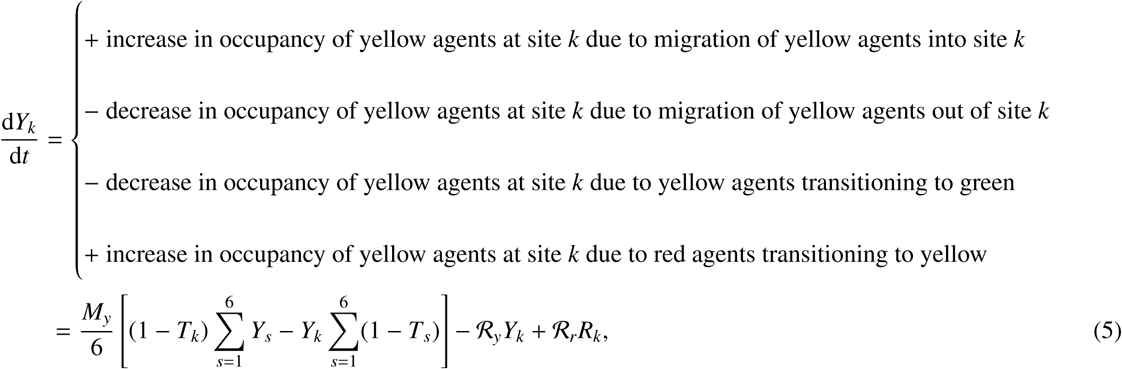

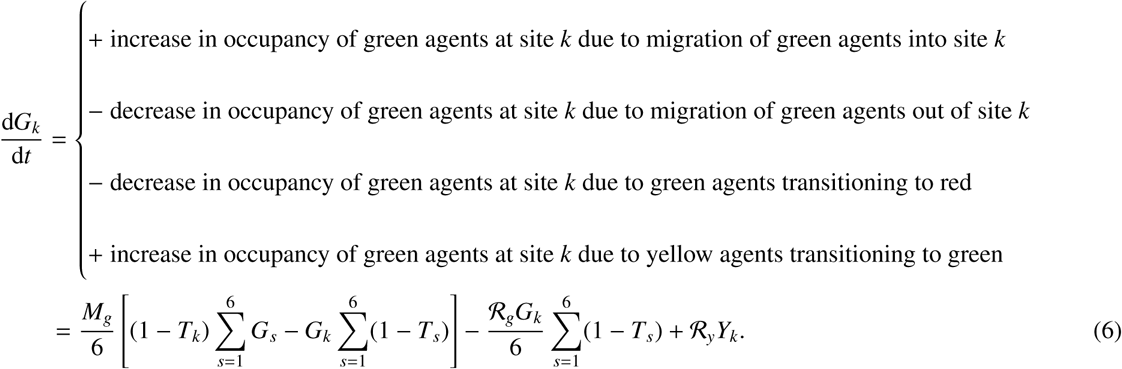

The index *s* in Equations (4)–(6) refers to the six nearest neighbouring lattice sites about site *k*, as indicated in Figure 1(d). This means that terms like 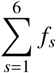 refer to summing *f* over the six nearest lattice sites. To be clear, this summation does not mean summing over the first six sites on the lattice.

All terms on the right of Equations (4)–(6) have simple physical interpretations. For example, the first term on the right of Equation (5) represents the increase in d*Y*_*k*_/d*t* as the sum of the probability that a yellow agent, located at a nearest neighbour lattice site, moves into site *k*. The summation over the six terms represents the fact that there are six such possible events, and the factor (1 - *T*_*k*_) incorporates exclusion effects since any possible event that would place an agent on an occupied site is aborted. The second term on the right of Equation (5) represents the decrease in d*Y*_*k*_/d*t* as the sum of the probability that a yellow agent, located at site *k*, moves out of site *k*. Again, there are six possible ways that a yellow agent at site *k* can leave that site, and the factor (1 - *T*_*s*_) is the probability that the target site is vacant. The third and fourth terms on the right of Equation (5) represent a yellow agent at site *k* transitioning into a red agent at site *k*, and where a red agent at site *k* transitioning into a yellow agent at site *k*, respectively. The terms on the right hand side of Equations (4) and (6) can be interpreted in a very similar way, with the exception that these discrete statements reflect the fact that the transition from green-to-red requires vacant space, and hence some of the terms proportional to the transition rates in Equations (4) and (6) are also proportional to (1 - *T*_*s*_).

To proceed to the continuum limit, each term in Equations (4)–(6) is expanded in truncated Taylor series about the central lattice site, and terms of 𝒪(Δ^3^) are neglected. These truncated Taylor series expansions are given in Appendix A. Considering the limit as Δ → 0, we identify *Y*_*k*_(*t*), *G*_*k*_(*t*), *R*_*k*_(*t*) and *T*_*k*_(*t*) with continuous functions, *Y*(*x, y, t*), *G*(*x, y, t*), *R*(*x, y, t*) and *T* (*x, y, t*), respectively, giving

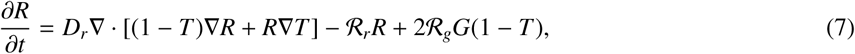

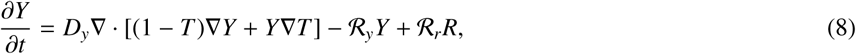

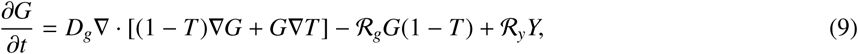

where 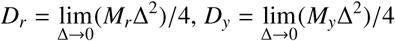 and 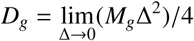 are the low-density diffusivity of the red, yellow and green subpopulations, respectively. To obtain a well-defined continuum limit we require that ℛ_*r*_ = 𝒪(Δ^2^), ℛ_*y*_ = 𝒪(Δ^2^) and ℛ_*g*_ = 𝒪(Δ^2^) as Δ → 0 [39]. It is worth noting that Equations (7)–(9) are analogous to the PDEs we studied in [16], however the key difference is that the diffusion terms in Equations (7)–(9) are nonlinear. The same type of nonlinearity arises when taking the continuum limit of exclusion processes that involve multiple species of subpopulations, as studied in [21].

Many kinds of two-dimensional assays in cell biology involve genuine two-dimensional movement of cells on substrates, however the initial condition is constructed in such a way that the density of cells depends only on one of the coordinates. Since this is the case in our experiments in Figures 1(a)–(b), we can simplify Equations (7)–(9) in the case where the density of cells depends only on the *x* coordinate, giving

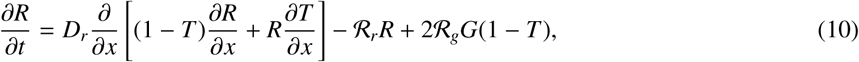

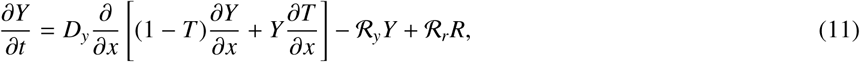

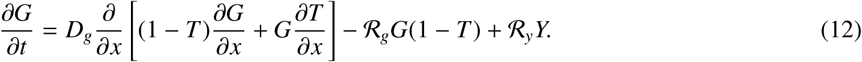

To solve Equations (10)–(12), we first discretise the spatial derivatives using a standard finite difference approximation with constant grid spacing, δ*x*. Backward Euler time stepping, with constant time steps of duration δ*t*, is used to step forward in time. Picard linearisation, with absolute convergence tolerance, ε, is used to solve the resulting system of nonlinear algebraic equations at each time step. The resulting system of linear equations is solved using the Thomas algorithm. All numerical results in this document correspond to δ*x* = 5 µm, δ*t* = 0.01 h and ε = 1 × 10^−5^. We find that these choices lead to grid-independent numerical results for the parameter combinations that we consider.

### 2.3. Applying the continuum and discrete models to mimic a scratch assay

The experimental procedure used to perform a scratch assay is outlined in Figure 2. To initiate the experiment, a population of cells, along with growth medium, is carefully placed into the wells of a tissue culture plate, as shown in Figure 2(a) [10, 16]. At this point we interpret the population of cells as being a spatially uniform monolayer, as indicated in Figure 2(b). A scratch is made in the monolayer of cells using a sharp tipped instrument [10]. The schematic in Figure 2(c) shows both the scratched region and the smaller imaged region. This schematic is important because it makes it clear that the images from this kind of assay show a small portion of the population of cells. In particular, the boundaries of the imaged region, shown in red in Figures 2(c)–(d), are not physical boundaries. Indeed, the spatially uniform population of cells extends far beyond the boundaries of the images. This means that whenever the density is below confluence, cells will migrate across the vertical red boundaries in Figure 2(c), and that the flux across these boundaries in the positive *x* direction will be equal in magnitude and opposite in direction to the flux in the negative *x* direction. Therefore, the appropriate mathematical boundary conditions along the vertical red boundaries in Figure 2(c) are zero net flux for each subpopulation. In our work we apply zero flux boundaries at both vertical boundaries in both the continuum and discrete formulations. In the discrete model this means aborting potential motility events that would place an agent across one of the vertical boundaries. In the continuum model this means enforcing zero net flux boundary conditions at both boundaries. Another important feature of the way that scratch assays are initialised is that while cells in the shaded regions in Figures 2(c)–(d) are free to move in two dimensions, the local density of cells in the shaded regions is constant at the beginning of the experiment, and this density is independent of the vertical location, *y*. Therefore, while cells are free to move in any direction, the density profiles depend only on the horizontal, *x*, direction. This means that we can model the migration and proliferation of cells in Figures 2(c)–(d) using a PDE where the independent variables are time, *t*, and horizontal coordinate, *x*, only. This approach to model a two-dimensional scratch assay using just one spatial variable is consistent with previous approaches [40].

**Figure 2:**
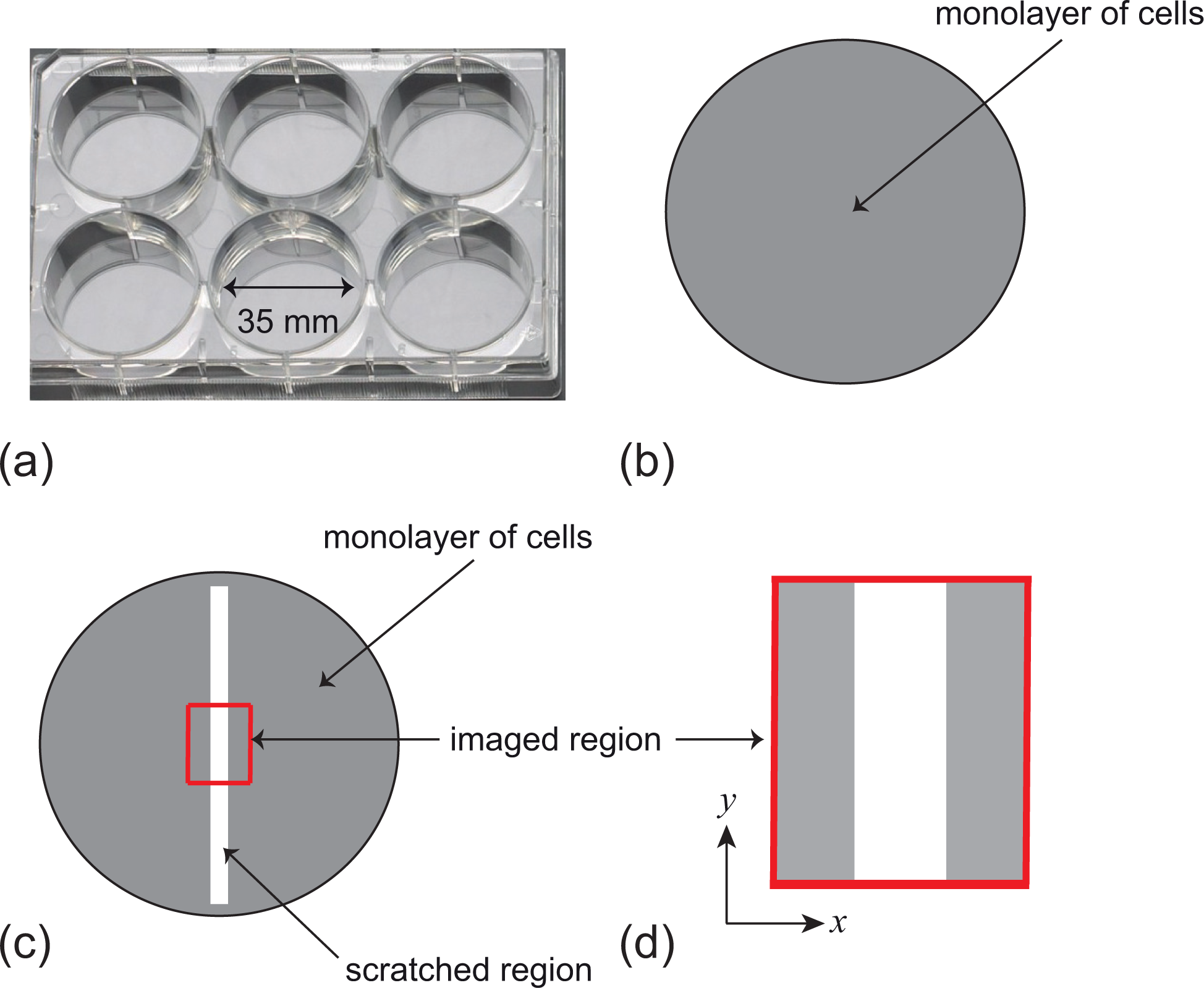
Experimental procedure explains the domain and boundary conditions for the stochastic simulations and continuum models. (a) Photograph of a six well tissue culture plate. Each well has a diameter of 35 mm. (b) Schematic showing cell monolayer (shaded) in the 35 mm diameter well. (c) Schematic showing the scratched region (white), the cell monolayer (shaded), and the smaller region that is imaged, outlined in red. (d) Magnified schematic of the imaged region showing the boundaries of the image, and the coordinate axes.

### 2.4. Parameterising the discrete model

To apply the discrete and continuum models to a particular experiment we need to specify both the initial condition and parameters in the model. The initial condition will be specified by using experimental images so that we initialise the location of agents in the discrete model to precisely match the location of cells at the beginning of the experiment. The initial condition for the continuum model will be specified by constructing the one-dimensional density profiles of the yellow, green and red subpopulations in the same experimental image at *t* = 0 h.

There are six rate parameters that need to be specified. To specify the transition rates we refer to Figure 1(c) in Haass et al. [12] which reports the average time that at least twenty individual 1205Lu melanoma cells spend in the G1, early S and the S/G2/M phases. These times are approximately 18 hours for the G1 phase, 4 hours for the early S phase, and 9 hours for the S/G2/M phases. Since these data are reported in terms of duration spent in each cell cycle phase, we convert these durations to rates by dividing ln 2 by the reported durations. Here the factor of ln 2 arises because the data reported by Haass et al. [12] corresponds to low density, freely cycling populations for which it is natural to assume that the population is growing exponentially, and doubling with each passage of the total cell cycle. Therefore, using these experimental data we set ℛ_*r*_ = ln 2/18 ≈ 0.04 /h, ℛ_*y*_ = ln 2/4 ≈ 0.17 /h and ℛ_*g*_ = ln 2/9 ≈ 0.08 /h. To specify the cell migration rates, we note that data presented by Haass et al. [12] show that the rate of cell migration in several melanoma cell lines is independent of the cell cycle phase. Therefore, we set *M*_*y*_ = *M*_*g*_ = *M*_*r*_ to reflect these experimental measurements. Furthermore, previous modelling studies suggest that the cell diffusivity for melanoma cell lines lies within the range 100-500 µm^2^/h [41]. Therefore, after some trial-and-error experimentation, we take an intermediate value of 400 µm^2^/h, and make use of the fact that a typical melanoma cell diameter is approximately 20 µm [41]. Together, these experimentally-motivated choices lead us to set *M*_*y*_ = *M*_*g*_ = *M*_*r*_ = 400 × 4/(20)^2^ = 4 /h. While all of the results presented here in the main document correspond to these experimentally-motivated choices, we present additional results in the Supplementary Material document where we systematically vary all six rate parameters to illustrate how the solutions of the model vary when we vary the rate parameters.

With these parameters we now demonstrate how to mimic the experimental images in Figures 3(a)–(b). Using ImageJ [42] we manually identify the (*x, y*) locations of all cells in Figure 3(a), and we visually classify them as being either red, yellow or green. These locations are mapped onto a hexagonal lattice, with Δ = 20 µm, as shown in Figure 3(c). A visual comparison of the images in Figure 3(a) and 3(c) confirms the accuracy of the mapping from continuous space to the hexagonal lattice. This process involves some approximation as the continuous position of each cell is mapped to the nearest lattice site on the hexagonal lattice. As we proceed through the list of cells, mapping each of them onto the lattice, we find that the closest lattice site for some particular cells is occasionally occupied. To deal with this scenario we simply map that cell to the next nearest site. Since our experiments are initialised at a relatively low density, we find that this situation arises infrequently. Once we have initialised the lattice, a single simulation is performed over *t* = 48 h, and the snapshot in Figure 3(d) shows the output. A comparison of the experimental image in Figure 3(b) and the simulation image in Figure 3(d) shows that the initially-vacant region has closed by this time, however since both the experiment and the simulation are stochastic we do not see the same match between the positions of the cells and agents by *t* = 48 h as we did at *t* = 0 h.

**Figure 3:**
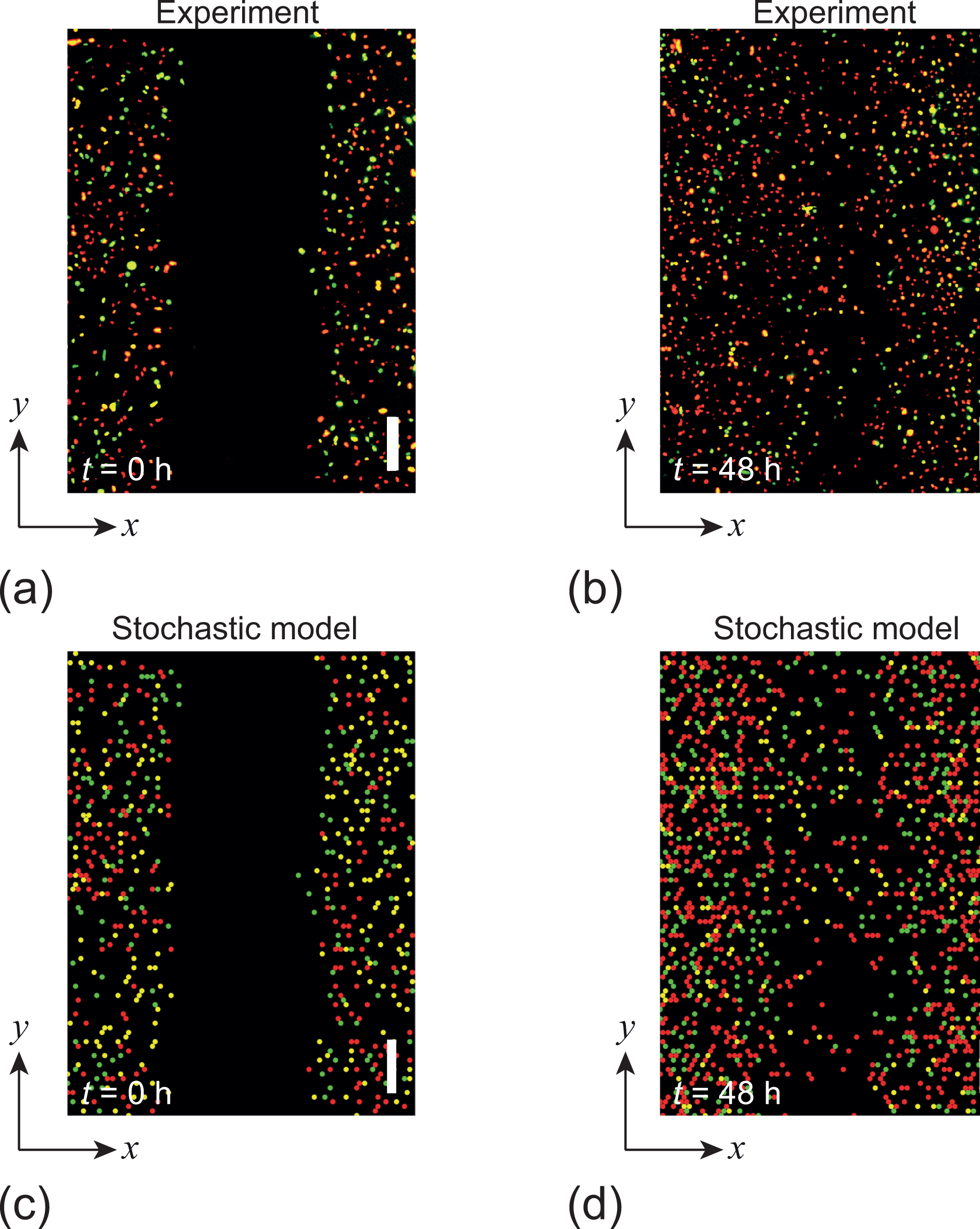
Visual comparison of experimental images and snapshots from the stochastic model. Results in (a)–(b) show images from a scratch assay using FUCCI technology in 1205Lu melanoma cells at *t* = 0 and *t* = 48 h, respectively. These are the same images shown previously in Figures 1(a)–(b). Images in (c)–(d) show snapshots from the stochastic model. Each scale bar corresponds to 200 µm. The snapshot in (d) corresponds to one realisation of the stochastic model with *M*_*y*_ = *M*_*g*_ = *M*_*r*_ = 4 /h, ℛ_r_ =0.04 /h, ℛ_y_ = 0.17 /h and ℛ_*g*_ = 0.08 /h. Zero net flux boundary conditions are imposed along all boundaries of the lattice, and the lattice spacing is set to Δ = 20 µm.

In addition to producing snapshots showing the position of agents from the stochastic model, we also extract density information from this kind of simulation. Results in Figures 4(a)–(c) show snapshots from the discrete model at *t* = 0, 24 and 48 h, respectively. This series of snapshots uses the same initial condition presented in Figure 3(c), which means that the initial condition is an accurate approximation of the initial condition in the experiment in Figure 3(a). Using these snapshots we construct density profiles, shown in Figures 4(a)–(c), by counting the number of agents, from each subpopulation, along each vertical column of sites. Dividing this number by the total number of sites per column produces a non-dimensional measure of the density, with maximum value of unity. These density profiles provide a way of quantifying how the density of each subpopulation varies in time and space. It is of interest to note that the density profiles in Figure 4(d) correspond to the placement of cells at the beginning of the experiment. These experiments are deliberately initialised with a very low density, which we can see is approximately 10-15% of the carrying capacity density. In this case, with Δ = 20 µm, the dimensional carrying capacity density of agents on the lattice is approximately 2.9 × 10^−3^ agents/µm^2^. It is important for our experiments that they are initialised at low density as this means that cells are relatively unaffected by contact inhibition and are therefore free to progress through the cell cycle. Another important consequence of working with such low density initial conditions is that this helps to avoid the situation where cells pile up in the vertical direction (which can happen when experiments are initialised at higher density). This is important because all mathematical models presented here make the implicit assumption that the population of cells forms a monolayer.

**Figure 4:**
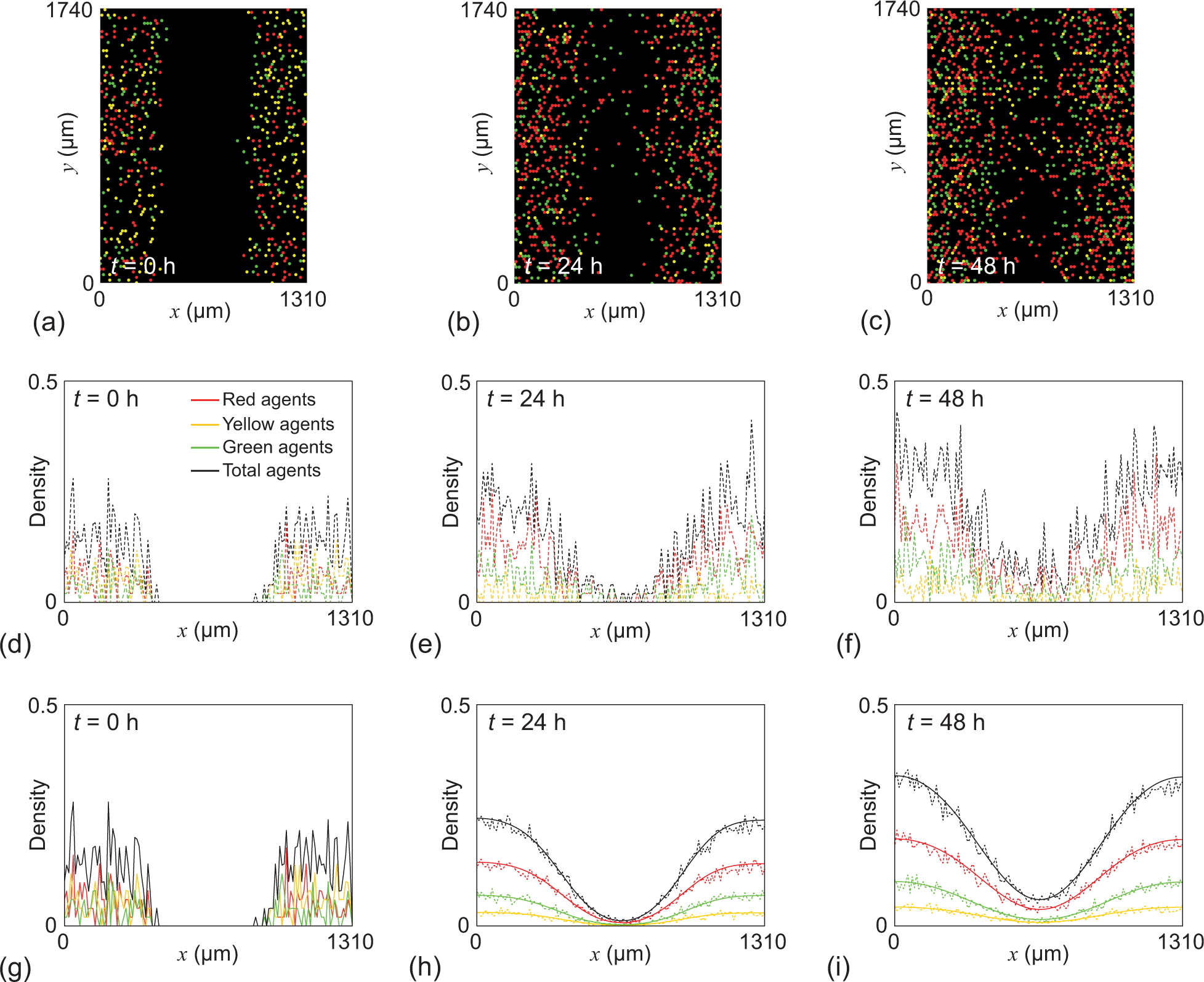
Extracting population-level density data from the stochastic model and comparing those results with the solution of the continuum limit PDE model. Results in (a)–(c) show a single stochastic simulation of a scratch assay using the same initial distribution of agents as in the experimental image in Figure 3(a). Snapshots are shown at *t* = 0, 24 and 48 h in (a)–(c), respectively. Results in (d)–(f) show density profiles obtained by taking the simulation results in (a)–(c) and counting the numbers of each agent type per column across the lattice, and dividing those numbers by the total number of sites per column. Results in (g)–(i) show averaged density profiles (dashed) obtained by repeating the stochastic simulations 20 times and averaging the resulting density profiles. Results in (g)–(i) also show solutions of Equations (10)–(12) (solid) at *t* = 0, 24 and 48 h, respectively. In all subfigures the red curves correspond to the density of the red subpopulation, the yellow curves correspond to the density of the yellow subpopulation, the green curves correspond to the density of the green subpopulation, and the black curves correspond to the total density. All PDE solutions are obtained on 0 < *x* < 1310 µm, with zero net flux boundary conditions at both boundaries. All results correspond to *M*_*y*_ = *M*_*g*_ = *M*_*r*_ = 4 /h, ℛ_*r*_ = 0.04 /h, ℛ_*y*_ = 0.17 /h, ℛ_*g*_ = 0.08 /h and Δ = 20 µm. Zero net flux boundary conditions are imposed along all boundaries of the lattice.

The data in Figures 4(d)–(f) from a single realisation of the stochastic model contains large fluctuations, and we can deal with this by repeating the stochastic simulations a number of times from the same initial condition, and averaging the resulting density profiles. Averaging each of these profiles produces relatively smoothed results, shown in Figures 4(g)–(i). To explore how well the continuum limit PDE description performs in this situation, we solve Equations (10)–(12) using the same initial condition in Figure 4(d), and superimpose these solutions in Figures 4(g)–(i). Comparing the continuous and stochastic density profiles in Figures 4(g)–(i) shows that the PDE description provides an accurate approximation of averaged data from the stochastic model. Perhaps the most striking benefit of working with the PDE approximation is that the solutions are very simple and fast to compute. For example, solving the PDEs to produce the plots in Figures 4(g)–(i) took less than two seconds of computational time on a single desktop computer. In contrast, performing 20 identically prepared realisations of the stochastic model required several minutes of computation. Therefore, the PDE description has the potential to reduce computational cost by at least an order of magnitude.

Now that we have demonstrated the continuum-discrete match for the scratch assay results in Figures 1(a)–(b), we consider whether the same quality of match holds over a longer timescale for a related problem where we alter the initial condition so that we see the formation of a single moving front of cells. First, we note that the kind of scratch assay geometry in Figure 3 leads to two opposingly directed fronts that meet and coalesce after a relatively short period of time. Therefore, this choice of initial condition and geometry does not lead to travelling wave-like solutions. To explore the potential for travelling wave-like solutions, we consider a slightly different initial condition that could correspond to a very wide scratch. In this related initial condition, we consider an initial population of cells that is confined near *x* = 0 with the remainder of the domain empty of any other cells. To formulate this initial condition we consider the population of cells towards *x* = 0 in Figure 1(a) on a much wider domain, 0 < *x* < 2500 µm, containing no other cells. Density data, obtained from 20 identically prepared realisations in Figures 5(a)–(d) show that this situation leads to a single front moving in the positive *x* direction. This single front is composed of a front of agents in the green subpopulation that is closely followed by a pulse of agents in the red and yellow subpopulations. Note that the timescale of the simulations in Figures 5(a)–(d) is much longer than the timescale of the simulations in Figure 4. Solutions of Equations (10)–(12) with the same initial condition are superimposed on the stochastic profiles in Figures 5(a)–(d), and comparing these PDE solutions with the averaged density profiles shows that the continuum limit PDE description provides an accurate description of averaged data from the stochastic model, even over these kinds of longer timescales required to see the formation of a travelling wave-like moving front.

**Figure 5:**
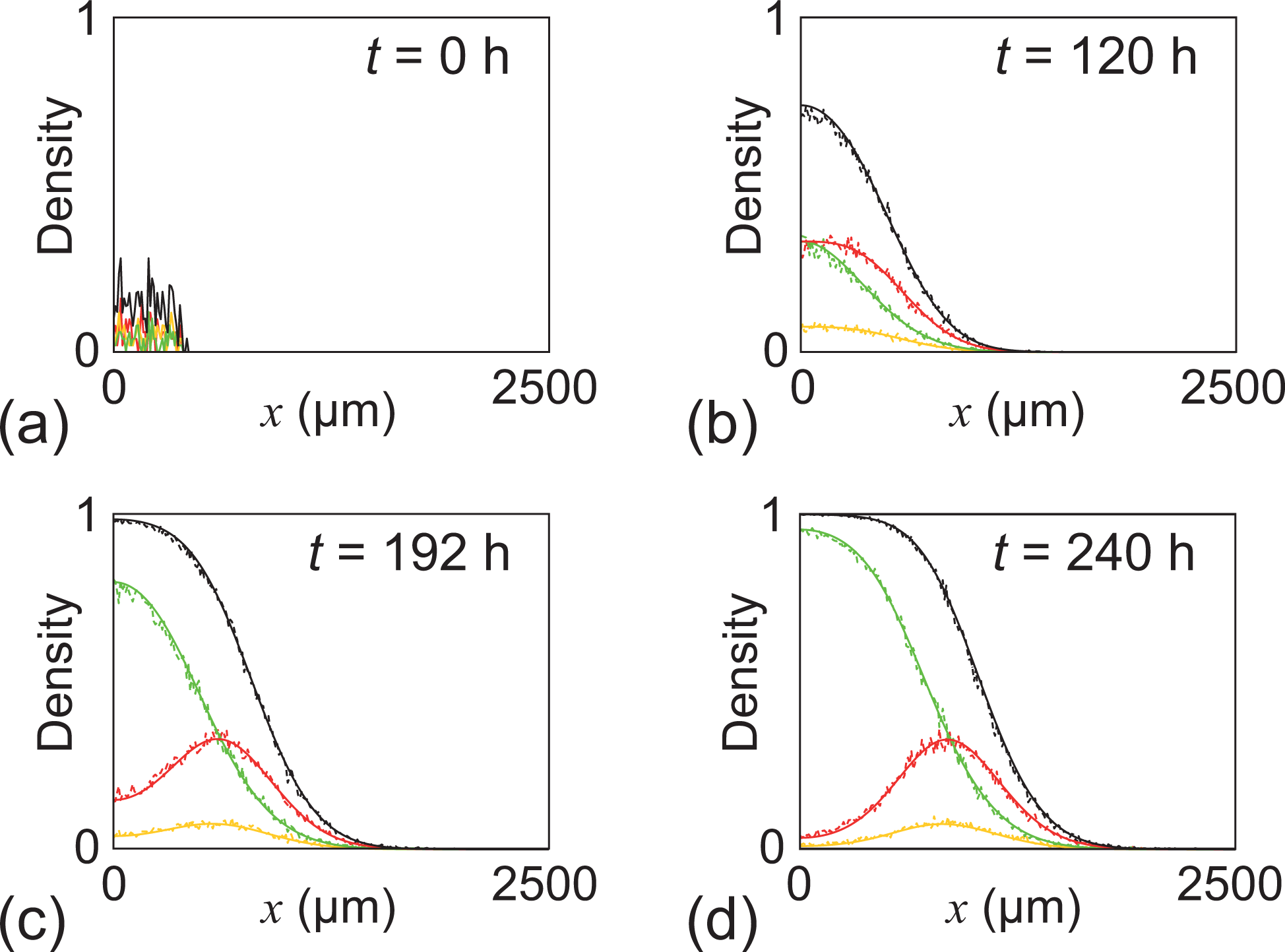
Longer stochastic simulations on a wider domain lead to travelling wave-type behaviour. Results in (a)–(d) show the density profiles (dashed) obtained by averaging results from 20 identically prepared realisations of the stochastic model. Results in (a)–(d) also show the solutions of Equations (10)–(12) (solid) for the same initial condition and boundary conditions. In all subfigures the red curves correspond to the density of the red subpopulation, the yellow curves correspond to the density of the yellow subpopulation, the green curves correspond to the density of the green subpopulation, and the black curves correspond to the total density. The domain is 0 < *x* < 2500 µm, and all simulations correspond to *M*_*y*_ = *M*_*g*_ = *M*_*r*_ = 4 /h, ℛ_*r*_ = 0.04 /h, ℛ_*y*_ = 0.17 /h and ℛ_*g*_ = 0.08 /h. Zero net flux boundary conditions are imposed along all boundaries of the lattice, and the lattice spacing is set to Δ = 20 µm.

All results showing the quality of the continuum-discrete match in Figures 4 and 5 correspond to one particular choice of parameters. This choice of parameters is useful because it directly reflects parameter estimates that are obtained from experimental observations [12, 41]. However, in addition to showing the quality of the continuumdiscrete match for this choice of parameters, we repeat the stochastic simulations and PDE solutions presented in Figure 5 for a range of parameter values and present those results in the Supplementary Material document. These additional results show that we still obtain a good quality continuum-discrete match, suggesting that the solution of Equations (10)–(12) provides an good approximate description of the stochastic process over a relatively broad range of biologically relevant parameter values.

## 3. Conclusion

In this work we present a new stochastic model of cell migration and proliferation that can be used to model two-dimensional cell migration assays incorporating fluorescent cell cycle indicators. This model involves treating the total population of cells as three interacting subpopulations: red agents model cells in the G1 phase of the cell cycle, yellow agents model cells in the early S phase, and green agents model cells in the S/G2/M phase. We explain how the stochastic model can be used to mimic cell biology assays using recently published data from a scratch assay with a melanoma cell line. Applying a mean field approximation, we derive a continuum limit PDE description of the stochastic model. Using repeated stochastic simulations, we show that the solution of the continuum limit PDE provides a good match to averaged data from the stochastic model.

The four main contributions of this work are to: (i) present a discrete model of cell migration that explicitly incorporates the cell cycle as highlighted by FUCCI technology; (ii) explain how to apply the discrete model to mimic a standard set of two-dimensional *in vitro* assays, called scratch assays; (iii) derive the mean-field continuum limit description of the stochastic model; and (iv) compare numerical solutions of the continuum limit model with averaged density data from repeated, identically prepared simulations of the stochastic model. In this work we do not consider calibrating the solution of the models to match experimental data. Instead, we simply adopt parameter estimates from previous studies [12, 41]. If we were to consider calibrating the models presented here to experimental data then it would be worthwhile taking great care to design the experiments so that we could learn as much as possible from the calibration exercise. The important new ingredient of the model is that it explicitly incorporates the cell cycle, and so it is useful to consider experiments initialised with a sufficiently low density of cells so that cells are relatively unaffected by contact inhibition and are freely cycling. Another key consideration is the width of the scratch used to initiate the scratch assay. Using a very narrow scratch would lead to rapid closure which would limit our ability to estimate the cell diffusivity. Instead, we would prefer to use a sufficiently wide scratch so that we have sufficient time to observe the movement of the red, green and yellow subpopulations into the wounded region. Perhaps the most important factor in calibrating the model to match experimental data is the consideration of fluctuations in the experiments. Data in Figure 4(d) show that averaging density information across one experimental image leads to very large fluctuations in density. One way of reducing these fluctuations is to consider performing a large number of identically-prepared experiments and averaging the density profiles across these experiments. However, when performing scratch assays it can be challenging to create scratches of the same width, and one way of overcoming this difficulty is to use a more controlled scratch assay, such as the IncuCyte ZOOM^™^ system that can be used to create reproducible scratch widths [11].

There are many ways that our present study could be extended. Here we assume that both the red-to-yellow transition and the yellow-to-green transition are unaffected by crowding whereas we assume that the green-to-red transition is affected by crowding since this transition requires the availability of space to accommodate a new daughter agent on the lattice. However, a more biologically realistic model might allow for each of the red-to-yellow, yellow-to-green and the green-to-red transitions to depend on the local density. This kind of generalisation, which could improve how the discrete model matches experimental observations at high density, could be incorporated into the discrete model by using more sophisticated measures of local density on the lattice [43]. Since our experiments focus on low-density situations where cells are freely cycling, we have not considered these kinds of extensions in the present work. However, we anticipate that such extensions will be important in our future work where we plan to consider similar experiments under higher density conditions. Another feature of our model that could warrant further exploration is the standard assumption that the time between events in the discrete model is exponentially distributed. Recently, Yates and colleagues have suggested that the assumption of exponentially distributed waiting times might not be appropriate when modelling cell proliferation [44]. Given additional information about the distribution of waiting times, it would be possible to modify the Gillespie algorithm to incorporate such information.

Another important aspect of our modelling that could be explored further stems from our observation that the numerical solution of the continuum limit PDE models in Figure 5 appears to approach constant speed, constant shape travelling wave solutions. In this work we have not formally analysed travelling wave solutions of Equations (10)–(12) since we focus on developing mathematical and computational tools that can be used to interpret images from cell biology experiments. Such experiments do not typically lead to travelling waves because of the choice of the initial condition, boundary conditions and experimental timescales considered. For example, typical experimental images in Figures 1(a)–(b) lead to two opposingly directed fronts that will meet before they have a chance to form a unidirectional travelling wave. In addition to this issue of geometry, the simulation results in Figure 5 suggest that even after a time period of 240 h the constant shape travelling wave profile is still developing for this choice of parameters. Nonetheless, it would still be interesting to examine travelling wave solutions of Equations (10)–(12) by introducing a change of coordinates, *z* = *x* - *ct*, where *c* > 0 is travelling wave speed solution moving in the positive *x* direction. This transformation leads to a system of six nonlinear first order ordinary differential equations that could be analysed in phase space using dynamical systems theory. The analysis of this system would be complicated by its high dimension and the nonlinear diffusion terms, and so we leave this for future consideration. Yet a further extension could be to apply the stochastic models and continuum limit PDE models to a different kind of experiment where the initial density of cells depends on both the *x* and *y* coordinates, such as a hole-closing problem [45]. In this case, the discrete model described here could be used directly to mimic this kind of experiment, but we would require the use of the two-dimensional continuum limit, given by Equations (7)–(9).

## 4. Appendix A: Taylor series expansions

The key step in deriving the continuum limit description is to use truncated Taylor series to express the occupancy of certain nearest neighbour lattice sites, depicted in Figure 1(d), to the central lattice site. For the lattice geometry and nomenclature in Figure 1(d), the following truncated Taylor series are useful

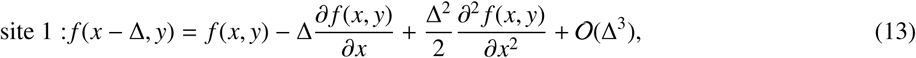

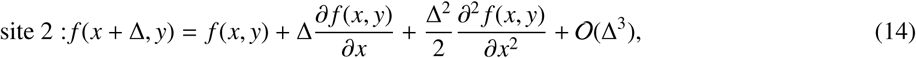

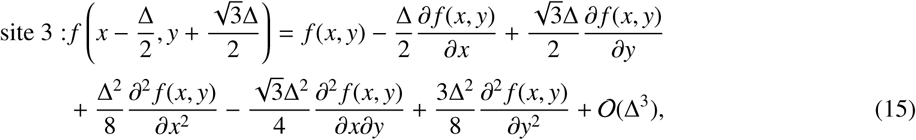

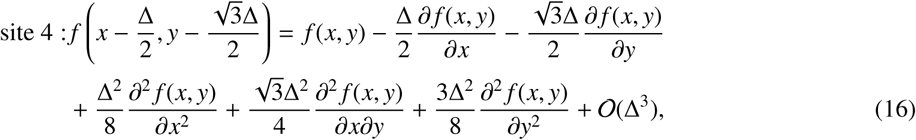

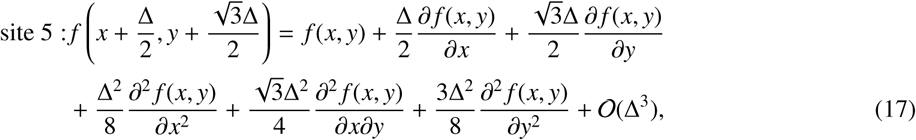

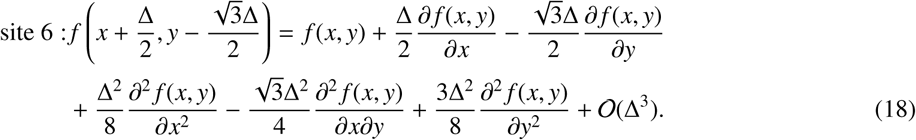

To derive the continuum limit PDEs we substitute the truncated Taylor series into Equations (4)–(6). Before doing so it is useful to identify certain key terms in these discrete conservation statements, such as 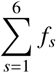, where *f*_*s*_ denotes the function, *f* (*x, y*), evaluated at the *s*^th^ nearest neighbour lattice site as shown in Figure 1(d). Summing Equations (13)–(18) allows us to write

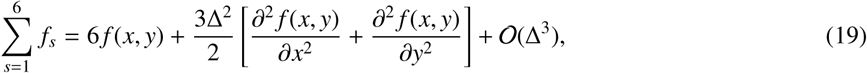

which is a useful result that we can use to simplify the algebraic expressions that we encounter when passing from Equations (4)–(6) to (7)–(9).

## Acknowledgements

This work is supported by the Australian Research Council (DP170100474). WJ is supported by a QUT Vice Chancellor’s Research Fellowship. JMR and TAT are both supported by the Australian Mathematical Sciences Institute (AMSI) through an AMSI Vacation research scholarship. NKH is a Cameron fellow of the Melanoma and Skin Cancer Research Institute, Australia, and is supported by the National Health and Medical Research Council (APP1084893). We thank four referees for their helpful comments and suggestions.

